# ProkBERT Family: Genomic Language Models for Microbiome Applications

**DOI:** 10.1101/2023.11.09.566411

**Authors:** Balázs Ligeti, István Szepesi-Nagy, Babett Bodnár, Noémi Ligeti-Nagy, János Juhász

**Author notes:** Correspondence: Balázs Ligeti.

## Abstract

Machine learning offers transformative capabilities in microbiology and microbiome analysis, deciphering intricate microbial interactions, predicting functionalities, and unveiling novel patterns in vast datasets. This enriches our comprehension of microbial ecosystems and their influence on health and disease. However, the integration of machine learning in these fields contends with issues like the scarcity of labeled datasets, the immense volume and complexity of microbial data, and the subtle interactions within microbial communities. Addressing these challenges, we introduce the ProkBERT model family. Built on transfer learning and self-supervised methodologies, ProkBERT models capitalize on the abundant available data, demonstrating adaptability across diverse scenarios. The models’ learned representations align with established biological understanding, shedding light on phylogenetic relationships. With the novel Local Context-Aware (LCA) tokenization, the ProkBERT family overcomes the context size limitations of traditional transformer models without sacrificing performance or the information rich local context. In bioinformatics tasks like promoter prediction and phage identification, ProkBERT models excel. For promoter predictions, the best performing model achieved an MCC of 0.74 for *E. coli* and 0.62 in mixed-species contexts. In phage identification, they all consistently outperformed tools like VirSorter2 and DeepVirFinder, registering an MCC of 0.85. Compact yet powerful, the ProkBERT models are efficient, generalizable, and swift. They cater to both supervised and unsupervised tasks, providing an accessible tool for the community. The models are available on GitHub and HuggingFace.

## 1 INTRODUCTION

Numerous tasks in bioinformatics involve classifying or labeling sequence data such as predicting genes (Sommer and Salzberg, 2021; Lukashin and Borodovsky, 1998; Delcher et al., 1999), annotating sequence features (Seemann, 2014; Tatusova et al., 2016; Meyer et al., 2019; Aziz et al., 2008), etc. A significant challenge in this field is deriving efficient vector representations from these sequences (Zhang et al., 2023). Classification tasks related to sequences – like classifying assembled contigs into MAGs (metagenome-assembled-genomes) or analyzing AMR-associated genes – are often addressed by initially categorizing the data into bins or using simple composition-based representations, such as k-mer frequency distributions. A common method involves converting sequences into a basic presence-absence vector, indicating whether a particular genome contains specific sequence features like mutations, motifs, or other patterns. However, a drawback of this method is that proximity in this representation space doesn’t always imply semantic similarity. Another prevalent representation uses hidden Markov models (Durbin et al., 1998), where the model parameters encapsulate the essential properties of the sequences. Yet, integrating such models with machine learning algorithms like support vector machines or random forests can be complex. Despite this, hidden Markov models have demonstrated their effectiveness in classification tasks and provide highest quality annotations (Cantalapiedra et al., 2021; Zdobnov and Apweiler, 2001).

Neural network-based representations have distinct advantages, primarily their compatibility with a wide range of machine-learning tools, including autoML and statistical frameworks. Past research has highlighted the effectiveness of neural network representations for sequences, with a variety of classification tasks addressed using networks such as CNNs and RNNs (Min et al., 2017). These networks have been employed in areas like motif discovery, gene-expression prediction (Kelley et al., 2018) splicing site recognition (Ji et al., 2021), and promoter identification, as detailed in several comprehensive reviews (Sapoval et al., 2022; Min et al., 2017; Zhang et al., 2023). However, convolutional neural networks face challenges, like the need for extensive labeled sequence data. They are also task-specific, limiting their applicability to other scenarios outside their training focus. A significant bottleneck in integrating neural networks into bioinformatics has been the scarcity of adequate labeled data. Yet, recent advancements in machine learning, inspired by breakthroughs in natural language processing, image analysis (Han et al., 2022), and protein structure prediction (Alipanahi et al., 2015; Jumper et al., 2021), have introduced new paradigms. Transformer-based architectures, especially large language models (Brown et al., 2020a; Raffel et al., 2020; Devlin et al., 2019), offer versatile representations – often termed “reusable” or “fundamental models”. Among the recent training approaches is the fine-tuning paradigm, which divides the training process into two phases: pretraining and fine-tuning. Pretraining demands vast amounts of self-labeled data, while fine-tuning can, in some instances, operate with minimal or even no examples.

In bioinformatics, there exists a paradoxical challenge. On one hand, there’s an abundance of sequence data available, especially in public repositories like the SRA (sequence read arhive). The volume of this data is expanding exponentially, and as sequencing and other data-producing technologies become more affordable, this growth trend is likely to persist. These data repositories are akin to hidden treasures. Yet, they remain under-analyzed and underprocessed. Researchers often focus primarily on specific mutations, neglecting other valuable aspects of the data. Conversely, while there’s an abundance of raw sequence data, there’s a scarcity of labeled data. The accompanying metadata is frequently limited, and given the high cost of experiments, only a handful of samples, typically ranging from 3–15, are available within a specific group or stratum. It’s also worth noting that labeling criteria can differ significantly across projects.

Recognizing these challenges, there is a compelling need for innovative methods that can harness the vast repositories of raw sequence data and navigate the complexity of labeling inconsistencies. It is in this context that our research contributes a novel solution. The development and application of our genomic language model family aims to address the mentioned issues, providing a robust, adaptable, and efficient tool for sequence classification.

While the concept of pretrained models isn’t new, several have emerged recently, such as DNABERT (Ji et al., 2021; Zhou et al., 2023), Nucleotide Transformer (Dalla-Torre et al., 2023), and LookingGlass (Hoarfrost et al., 2022). However, a common limitation among these methods is their primary focus on human sequences or their restricted context size.

In the pretraining phase, the objective is to derive a general representation that captures the semantic relationships between objects, which in this context means obtaining a nuanced representation of sequence data. Typically, achieving this requires billions of samples, yet the volume of available sequence data far surpasses this number. We trained our genomic language model on an extensive corpus of available sequence data, encompassing bacteria, archaea, viruses, and fungi. Subsequently, we fine-tuned our models to tackle specific classification tasks, including the recognition of promoters and phages.

The ProkBERT family encompasses a series of models tailored to meet the intricate demands of microbial sequence classification, analysis, and visualization. The versatility of the ProkBERT models is manifested through their diverse applications:

1. Zero-shot learning: This approach allows for clustering of sequences by leveraging the embeddings directly produced by the model, eliminating the necessity for explicit fine-tuning.
2. Sequence classification: ProkBERT models can be seamlessly fine-tuned, whether for token-specific or comprehensive sequence-based classification tasks.

With these capabilities, the ProkBERT family aims to bridge the current gaps in the field, offering a robust toolset for diverse bioinformatics challenges.

## 2 MATERIALS AND METHODS

In this study, we used the transfer-learning paradigm for sequence classification based on transformer-based architectures. The first phase involves pretraining on a large amount of sequence data, allowing the model to learn general sequence patterns. Once this foundation is established, we move to the fine-tuning phase where the model is adapted to specific tasks or datasets. The following sections provide a step-by-step description of our methods, from preparing raw sequence data to the specifics of both pretraining and fine-tuning. Figure 1 illustrates the training process.

**Figure 1.**
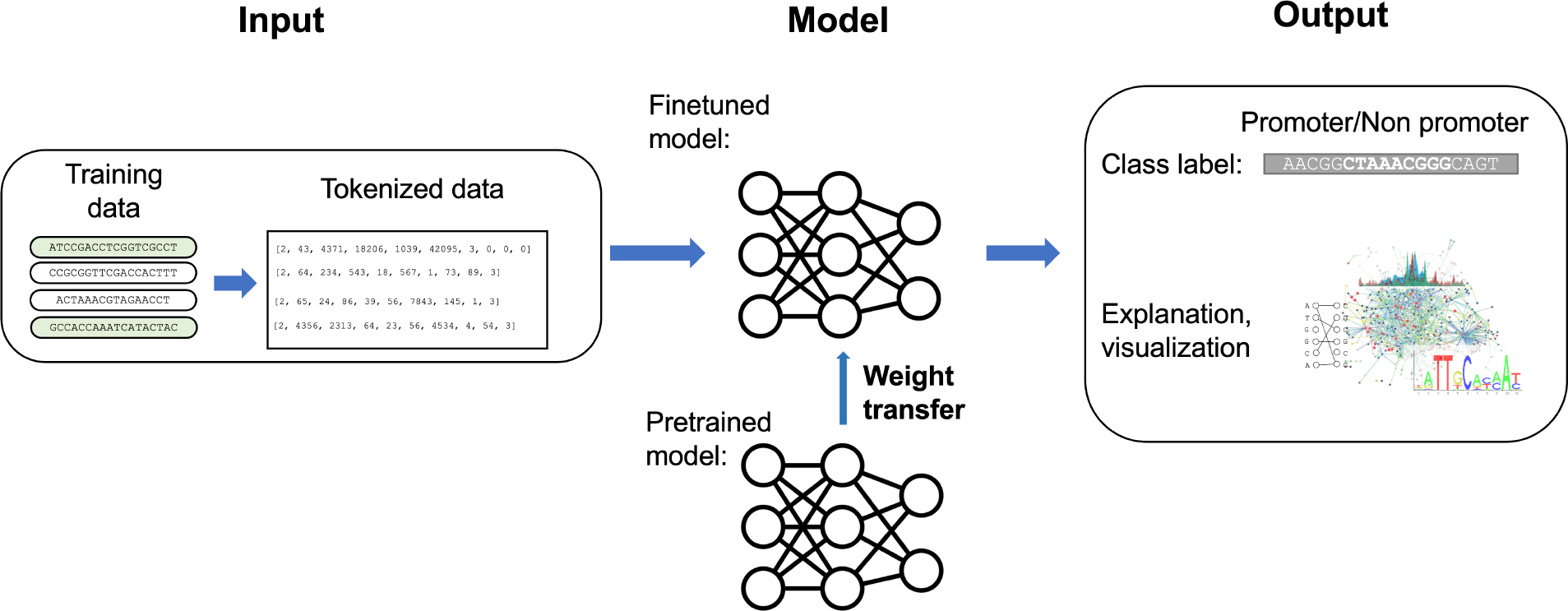
A schematic overview of the training process. Starting with raw sequence data, it undergoes preprocessing and vectorization. The model is then fine-tuned, beginning with weights from a pretrained model, to address the specific classification task. The output showcases classified sequences or tokens, predicted labels, scores, and a visualization highlighting underlying sequence patterns and explanations.

In the development of the ProkBERT family, the initial step involves pretraining the model on a vast corpus of data. During this pretraining phase, the model aims to tackle the Masked Language Modeling task. In this task, specific portions of the sequence are masked, and the model’s objective is to predict these masked sections, optimizing the likelihood of the missing parts using cross-entropy as the loss function. The model typically receives input in the form of a vectorized representation of the sequence. A notable constraint of standard transformers is their limited input size. Though various solutions have been suggested to address this limitation, the maximum token size is typically restricted to 0–4kb, significantly smaller than the average bacterial genome, but much larger than an average gene.

Fine-tuning nucleotide sequences is a technique used to adapt pre-trained models to specialized tasks or specific datasets. The first step involves segmenting raw sequences into chunks, usually ranging from 0–1kb in size, to optimize the model’s learning capability (Pan and Yang, 2009). Using weights from a pre-trained model, the system benefits from the knowledge obtained from comprehensive training on extensive datasets (Vaswani et al., 2017; Devlin et al., 2019). This initialization helps in quicker convergence and improved performance. After this initialization, the model undergoes training on the desired dataset, adjusting to its specific patterns and details. The outcome of this procedure allows the model to produce labeled sequences or tokens, which can be used for various annotation or prediction purposes (Brown et al., 2020b).

### 2.1 Sequence data

#### 2.1.1 Sequence segmentation and tokenization

The first step is processing the sequence data. While there are many parallels between sequence data processing and natural language processing, drawing direct analogies can be challenging. For instance, determining what constitutes a ‘sentence’ in the realm of nucleotide and protein sequences doesn’t have a direct counterpart in natural language. Additionally, the input size for neural networks is inherently limited. Figure 2 illustrates the strategy employed to vectorize the sequences.

**Figure 2.**
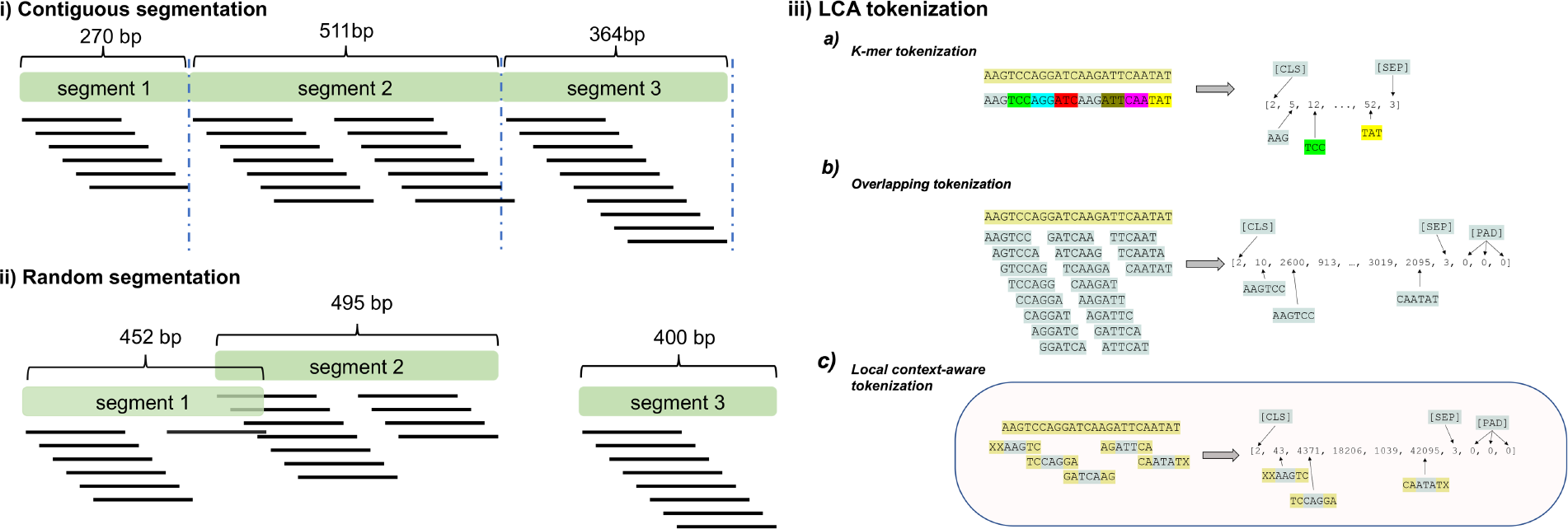
Preprocessing of sequences. The sequences are initially segmented into chunks ranging between 0–1kb. Two segmentation strategies can be employed: i) *Contiguous segmentation* where the sequence is split into non-overlapping parts; ii) *Random segmentation* where segments of varying lengths are randomly sampled from the original contig. The third part (iii) outlines the tokenization process of the segments: a) Splitting segments into non-overlapping tokens; b) Creating maximally overlapping k-mers; c) Generating partially overlapping k-mers by shifting with a fixed size.

Initially, the input sequence is segmented into smaller chunks. We employed two approaches for this:

1. Contiguous sampling, where contigs are divided into multiple non-overlapping segments; and
2. Random sampling, which involves fragmenting the input sequence into various segments at random.

Following segmentation, the next phase is encoding the sequence into a simpler vector format. The primary question revolves around defining the fundamental building block for a token. Various solutions have been suggested, the most widely strategy is applying one-hot-encoding (Sapoval et al., 2022), but DNA-BERT (Ji et al., 2021) applies the maximal overlapping k-mer strategy, meanwhile others relies on nucleotide level mapping (Dalla-Torre et al., 2023).

This phase is termed tokenization. We introduce a method termed *Local Context Aware* tokenization (LCA), where individual elements consist of overlapping k-mers. Two principal parameters dominate this approach: k-mer size and shift. For *k* = 1, the tokenization resorts to a basic character-based approach, with a typical example illustrated in Figure 2.

Employing overlapping k-mers can lead to enhanced classification performance. A greater shift value allows the model to use a broader context while reducing computational demands, while having the information-rich local context as well.

In this study, we propose models with a k-mer size of 6 (termed `ProkBERT-mini`), k-mer size of 1 (dubbed `ProkBERT-mini-c`), and a variant supporting a larger context window, named `ProkBERT-mini-long`, which relies on a k-mer size of 6 with a shift = 2.

#### 2.1.2 Training data

The dataset was retrieved from the NCBI RefSeq database (O’Leary et al., 2016; Li et al., 2021) on January 6th, 2023. It included reference or representative genomes from bacteria, viruses, archaea, and fungi. After filtering, the sequence database consisted of 976,878 unique contigs derived from 17,178 assemblies. These assemblies represent 3,882 distinct genera, amounting to approximately 0.18 petabase pairs. The segment databases was created by sampling fixed lengths of [256, 512, 1024] or, in other instances, variable lengths aiming for an approximate coverage of 1.

Tokenization was performed using various k-mer sizes and shift parameters. The compiled database was then stored in the Hierarchical Data Format (HDF). Collectively, the training database held roughly 200 billion tokens for each segmented dataset.

For transparency and further research, all training data is available at zenodo 10.5281/zenodo.10057832.

### 2.2 Pretraining and learning sequence representations

#### 2.2.1 Transformer model selection and parameters

In our study, we employed the MegatronBertForMaskedLM model (Shoeybi et al., 2019), a variant of the BERT architecture (Devlin et al., 2019), optimized for large-scale training. The key attributes of our models can be seen in Table 1.

**Table 1.** A comprehensive overview of model parameters across varied configurations.

|  | mini | mini-c | mini-long |
| --- | --- | --- | --- |
| parameters | 20,6 m | 24,9 m | 26,6 m |
| tokenizer | 6-mer, shift=1 | 1-mer | 6-mer, shift=2 |
| layers | 6 | 6 | 6 |
| attention heads | 6 | 6 | 6 |
| max. context size (bp) | 1027 nt | 1022 nt | 4096 nt |
| training data | 206.65 billion | 206.65 billion | 206.65 billion |

The `mini` and `mini-long` models has a vocabulary of 4,101 k-mers, while the `mini-c` employs 9 (comprising special tokens and nucleotides). We applied a learnable relative key-value positional embedding, mapping input vectors to a 384-dimensional space. The maximum supported lengths are 1024bp and 2048 bp, respectively. Intermediate layers of the encoder use the GELU activation function, expanding input dimensions to a 3,072-dimensional space before compressing back to 384 dimensions. The Masked Language Modeling head was designed with a decoder mapping from 384 to 4,101 dimensions. To ensure efficient parallel computations, the entire architecture was encapsulated within a DataParallel wrapper, optimizing GPU utilization. For implementation, we relied on the PyTorch version 2.01 framework and the Hugging Face library version 4.33.2.

#### 2.2.2 Training process

##### 2.2.2.1 Masked Language Modeling objective modifications

While Masked Language Modeling (MLM) acts as the primary pre-training objective for BERT models (Bidirectional Encoder Representations from Transformers) as established by Devlin et al. (2019), our implementation has slight variations. In the traditional BERT approach, a certain percentage of input tokens are randomly masked, and the model predicts these based on their context. Typically, about 15% of tokens undergo masking. However, due to our usage of overlapping k-mers, masking becomes more intricate. If a k-mer of size *k* = 6 is masked, we need to ensure at least six tokens are also masked to prevent trivial restoration from context and locality.

For an input sequence of tokens **x** and a binary mask vector **m** – where 1 indicates a masked token and 0 indicates an unmasked token – the model outputs predicted vectors **y**. As for the noise application on masked tokens, probabilities denotes the predicted probability nd *p*_3_ define different noise strategies. In our model, when a token is masked, it is substituted with the special `[MASK]` token with a probability of *p*_1_. Alternatively, with a probability *p*_2_, it can be replaced with a random k-mer from our vocabulary. Lastly, there’s a *p*_3_ chance that the masked k-mer will remain as it is. Conventionally, these probabilities are set at 0.8, 0.1, and 0.1, respectively.

The MLM objective aims to minimize the negative log likelihood over all masked positions, as described by the equation:

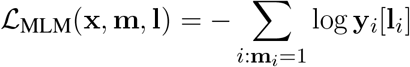

Where **y**_*i*_[**l**_*i*_] denotes the predicted probability of the true label **l**_*i*_ for the masked position *i*. This objective, coupled with the noise injection strategy, ensures that the model learns bidirectional representations, thus becomes capable of understanding and generating contextually relevant tokens.

When dealing with overlapping k-mers, simple token masking becomes insufficient. If a single k-mer token is masked, all overlapping k-mers related to that token must also be masked. This is crucial because when a k-mer is not masked and subsequently restored, it might inadvertently provide contextual information about its neighbors. Such a situation would enable the trivial restoration of adjacent masked k-mers. In essence, one unmasked k-mer could potentially “leak” enough information to unmask its neighboring tokens. For examples, as presented in Figure 2/iii (*Overlapping tokeniization*), if only the second token ‘AGTCCA’ is masked, it can be fully restored from its neighbouring tokens: ‘AAGTCC’ and ‘GTCCAG’.

This overlapping nature of k-mers posed unique challenges. As a result, we had to dynamically adjust the MLM parameters and the lengths of sequence segments during the pretraining phase. Additionally, when multiple contiguous k-mers were masked together, the probability associated with the MLM had to be recalibrated. This was necessary to ensure that the actual proportion of the sequence being masked was consistent with our intended masking ratio.

##### 2.2.2.2 Training phases and configuration

Initially, we employed parameters that allowed complete sequence restoration (k-mer of *k* = 6) by masking only five continuous tokens (with *p*_1_ = 0.9) and manipulating 15% of the tokens. Once a loss threshold of 1 was attained, the MLM parameters were adjusted to heighten the masking complexity. We implemented various masking lengths, such as 2 nucleotides for k-mer of *k* = 6 and 2 characters for *k* = 1. Training data in the first phase had a fixed length of 128nt segments. The succeeding phase used variable-length datasets: with a probability of 0.5 a full-length segments, and with a probability of 0.5 a segment between 30-512bp was selected into the the batch. The termination criterion for training was no further improvement or performance decrease, in both the MLM and promoter tasks. Models underwent training for roughly one batch each. We opted for batch sizes that spanned around 0.5-2 million bp sequences. Computations were executed on HPC-VEGA and Komondor platforms with Nvidia-A100 GPUs, leveraging slurm, pytorch distributed, and multiple GPU nodes.

#### 2.2.3 Evaluating the pretrained model

We evaluted the masking performance of the models using the ESKAPE pathogens, namely *Enterococcus faecium* (GCF 009734005.1), *Staphylococcus aureus* (GCF 000013425.1), *Klebsiella pneumoniae* (GCF 000240185.1), *Acinetobacter baumannii* (GCF 008632635.1), *Pseudomonas aeruginosa PAO1* (GCF 000006765.1) and *Escherichia coli str. K-12* (GCF 000005845.2), because of their high clinical importance. First we investigated how the genomic structure is reflected in the embeddings, on different sequence features (i.e. CDS, intergenic, pseudo-genes, etc.). Next we measured how well the models can perform in masking.

#### 2.2.4 Analysis of encoder outputs

In deep learning, an encoder typically processes input data (such as a sequence of tokens) and produces a dense vector representation for each token. These dense vectors, often referred to as embeddings or encoded vectors, capture the semantic information of the input tokens.

Given an input sequence *S* with *T* tokens, i.e.,

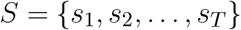

the encoder produces a sequence of vectors:

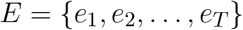

where *e*_*i*_ represents the embedded vector for the token *s*_*i*_.

In case of multiple inputs or batches, if we have a batch of size *B* with each sequence containing *T* tokens, the encoder’s output would be a 3D tensor of shape (*B, T, D*) where *D* is the dimensionality of the embeddings.

Once we have the encoded vectors, there are several ways to aggregate or pool them to get a single representation for the entire sequence. Here are some common pooling methods:

- **Mean Pooling:** Average the vectors: 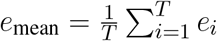.
- **Sum Pooling:** Sum the vectors: 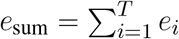.
- **Max Pooling:** Max value per dimension: 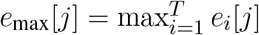.
- **Min Pooling:** Min value per dimension: 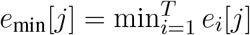.

For batches, these pooling operations are applied independently for each input sequence in the batch. The provided NCBI annotations were preprocessed and extended. Intergenic regions were defined as non-annotated genomic features with respect to the strand. We retained the CDS, intergenic, pseudo-genes, ncRNA features, while the rare or infrequently used features (such as riboswitch, binding_site, tmRNA, etc.) were excluded from the analysis. This was followed by sampling segments of various lengths from each genomic region. We sampled a maximum of 2000 sequence features from each contig, considering the strand, to evaluate strand-specific biases as well.

Then, we randomly corrupted a segment 10,000 times, i.e., a character was replaced with ‘*’ and tokens containing ‘*’ were mapped to the [MASK] token as illustrated on Figure 3. The sampled segment database is available at Zenodo 10.5281/zenodo.10057832.

**Figure 3.**
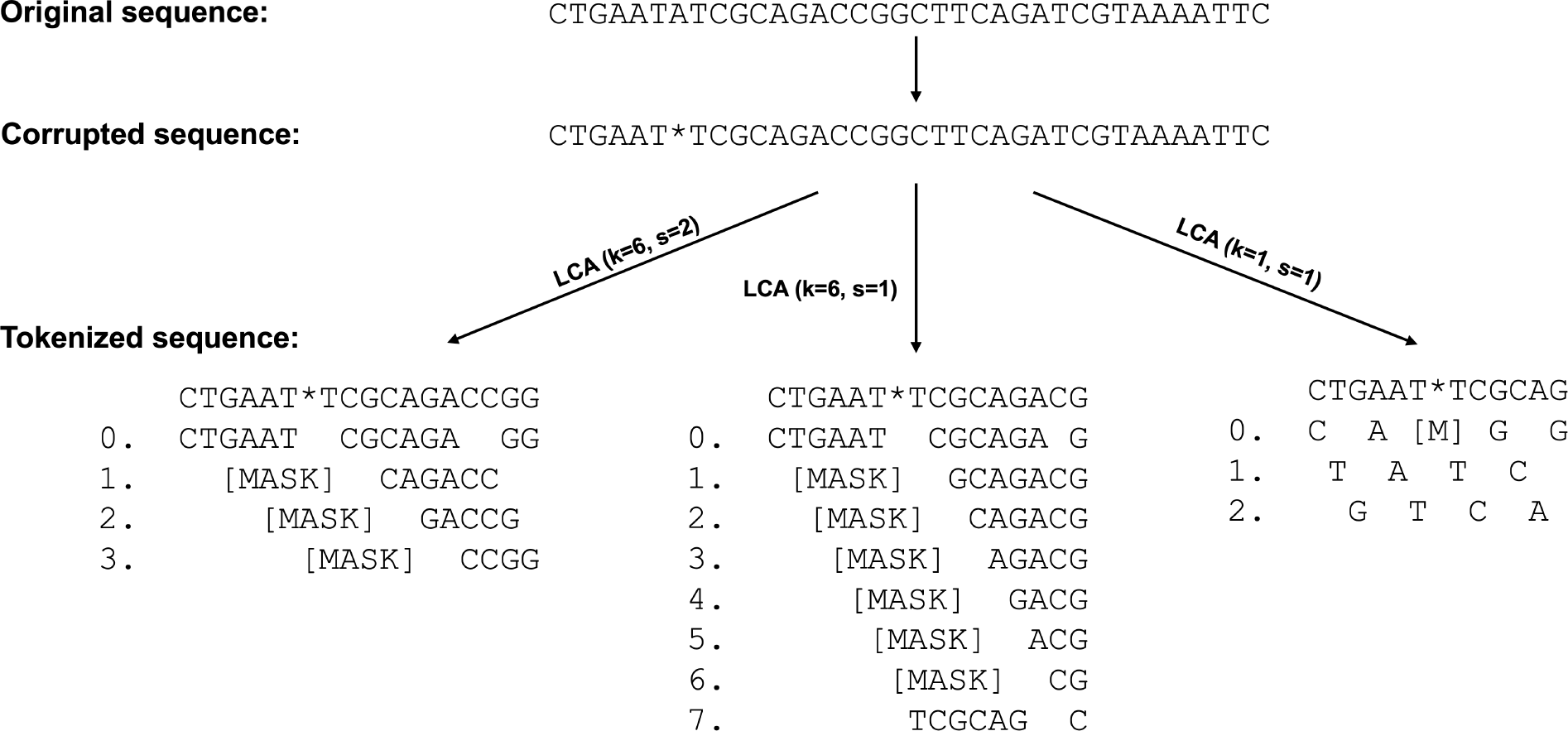
LCA Tokenization and corrupted sequence restoration. The figure illustrates how the corruption of a character at sequence level affects the initial vector representations of the seqment with respect to the different tokenization methods. The 7th nucleotide is unknown or masked, and those k-mers that overlap with that position are masked. As a result, when *k* = 6 and *s* = 2 not only the 7th character is hidden, but the 8th as well.

### 2.3 Application I: Bacterial promoter prediction

#### 2.3.1 Dataset overview

The first task our models were evaluated on involved distinguishing between promoter and non-promoter sequences in bacteria. A sequence is labeled ‘1’ if identified as a promoter and ‘0’ otherwise.

Our promoter database primarily built on Prokaryotic Promoter Database (PPD, Su et al., 2021), containing experimentally validated promoter sequences from 75 organisms.

For validation, we referred to Cassiano and Silva-Rocha (2020)’s dataset comprising *E. coli* sigma s70 sequences. The positive, well-recognized samples came from Regulon DB (Santos-Zavaleta et al., 2019). Cassiano and Silva-Rocha (2020) evaluated various tools, using an experimentally validated *E. coli* K-12 promoter set dependent on sigma 70, sourced from Regulon DB 10.5 (Santos-Zavaleta et al., 2019). Given the extensive documentation of sigma 70-dependent promoters in bacteria, only these were taken into consideration. They used a positive dataset of 865 high-evidence sequences from Regulon DB and a negative set of 1,000 sequences mimicking the nucleotide distribution of the natural sequences. Each sequence was 81 bp long, ensuring compatibility with most tools’ input prerequisites, particularly around the putative TSS region interval [*−*60, +20].

Our positive dataset encompasses promoter sequences from various species, predominantly found on both chromosomes and plasmids. We rigorously ensured no overlap existed within the promoter datasets.

The alternative test set consists of promoters from multiple species. The positive promoters were randomly divided into portions of 0.8, 0.1 and 0.1, as shown in Figure 4. We then evaluated the models using this dataset as well.

**Figure 4.**
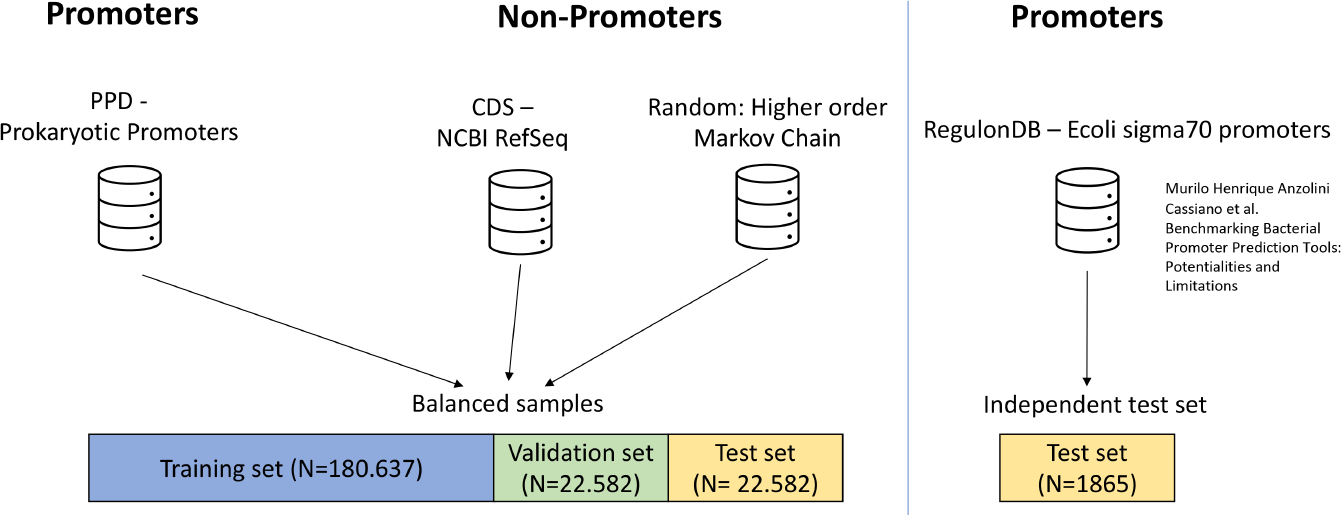
Promoter dataset schematic. A visual representation of the source and distribution of sequences used in the study, segregating prokaryotic promoters from non-promoters, with an additional independent test set based on *E.coli* sigma70 promoters.

To curate a comprehensive negative training set, we employed three strategies:

1. Using non-promoter sequences (CDS – Coding Sequences).
2. Random sequences generated with a 3rd-order Markov chain.
3. Pure random sequences.

The distribution of this composite dataset was 40% CDS, 40% Markov-derived random sequences, and 20% pure random sequences. Subsequent to the amalgamation, we partitioned the dataset into training, testing, and validation subsets.

A 3rd-order Markov chain predicts the next nucleotide in a sequence based on the states of the previous three nucleotides. Formally, the probability of observing a nucleotide *x*_*i*_ given the nucleotides at positions *x*_*i−* 3_, *x*_*i−*2_, and *x*_*i−*1_ is:

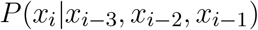

For DNA sequences, this yields 4^4^ = 256 possible nucleotide combinations. Such higher-order modeling can more effectively capture intricate sequence patterns and dependencies than lower-order models (Durbin et al., 1998). However, estimating transition probabilities requires extensive data due to the increased number of states (Koski and Noble, 2001). We determined these probabilities using promoter sequences, to which we added the reverse complement of each promoter. Subsequently, random promoter sequences were generated using these models. The CDS sequences were extracted from the genomic sequences of contigs, based on annotations from NCBI.

Figure 4 shows a schema of the promoter database.

#### 2.3.2 Training for promoter prediction

We employed a fine-tuning paradigm to evaluate our model. Our proposed binary classification model extends the Megatron BERT architecture (Shoeybi et al., 2019), tailored specifically for binary classification tasks. Let **X** represent the sequence of input embeddings, with *f*_BERT_(**X**) denoting the transformation by Megatron BERT. Given an input sequence of length *T*, this model transforms **X** into a sequence output **S** with dimensions *T ×* hidden size, where **S** = *f*_BERT_(**X**). Unlike the conventional BERT model, which classifies sequences based on the special `[CLS]` token representing the ‘sentence’, our approach emphasizes integrating representations of all tokens using a weighting scheme.

To obtain a fixed-size representation from the variable-length sequence **S**, we devised a weighting mechanism. The sequence **S** undergoes a transformation through a linear layer to yield a sequence of weights **W**:

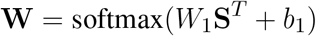

Here, *W*_1_ is a matrix sized hidden size *×*1 and *b*_1_ is a bias term. The softmax operation ensures **W** forms a valid probability distribution over sequence positions. The model then computes a weighted sum of the sequence representations:

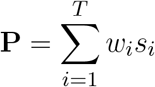

Where *w*_*i*_ and *s*_*i*_ represent the weight and the sequence representation at the *i*^*th*^ position, respectively.

Subsequently, **P** is processed by a dropout layer with a probability of hidden dropout prob to produce **P**^*′*^. This results in the final classification logits **L**.

Datasets, comprising training, validation, and testing subsets, were appropriately tokenized and adapted for ProkBERT processing. For optimization, the AdamW variant was chosen with parameters *α∈* {0.0001, 0.0004, 0.0008 }, *β*_1_ = 0.95, *β*_2_ = 0.98, and *ϵ* = 5 *×* 10^*−*5^. A linear learning rate scheduler with warmup was utilized. The model underwent training for two epochs, with a batch size of 128 per GPU (NVIDIA A100-40GB GPUs) using the pytorch data distributed framework (nvcc). Additional configurations included a weight decay of 0.01.

### 2.4 Application II: Phage sequence analysis

Bacteriophages have a significant role in the microbiome, influencing host dynamics and serving as essential agents for horizontal gene transfer (De la Cruz and Davies, 2000). Through this mechanism, they aid in the transfer of antibiotic resistance and virulence genes, promoting evolutionary processes. Understanding the diversity of phages is crucial for tackling challenges like climate change and diseases (Jansson and Wu, 2023). These phages exhibit distinct patterns in both healthy and diseased microbiomes (Yang et al., 2023). The correlation between the human virome and various health conditions, such as cancer, inflammatory bowel diseases, and diabetes, has been documented (Fernandes et al., 2019; Liang et al., 2020; Zuo et al., 2022; Han et al., 2018; Nakatsu et al., 2018; Zhao et al., 2017). However, deeper research is needed to discern causality and their impact on microbial and host biological processes.

Despite the abundance of phages (Bai et al., 2022a), accurately quantifying and characterizing them remains a challenge. One primary limitation is the restricted number of viral sequences in databases like NCBI RefSeq. Additionally, the categorization of viral taxonomy is still a topic of discussion (Walker et al., 2022). Though there have been recent efforts to expand databases (Camargo et al., 2023; Zhang et al., 2022), the overall understanding of viral diversity is still not complete (Yan et al., 2023). We have assembled a unique phage sequence database using recently published genomic data.

Another challenge is the life cycle of phages; temperate phages might integrate their genomes into bacterial chromosomes and are often annotated as bacterial genomes, leading to potential misidentification. Current databases also show biases towards certain genera (Schackart III et al., 2023), which can skew benchmarking and the evaluation of different methods. To address this, we used a balanced benchmarking approach, ensuring each viral group corresponds to their predicted host genus, minimizing bias. We also compared viral genomes to their respective hosts, a more demanding classification task, such as distinguishing a *Salmonella* phage from its host genome compared to marine *cyanobacteria*. For our study, we selected a specific number of phages for testing, ensuring there is no overlap between training and testing sets at the species level.

#### 2.4.1 Phage dataset description

To train and assess our prediction models, we assembled a comprehensive phage sequence database from diverse sources. As of 9th July, 2023, we procured viral sequences and annotations from the RefSeq database (O’Leary et al., 2016; Li et al., 2021). By isolating entries labeled ‘phage’, we obtained 6,075 contigs. Our database was further enriched with the inclusion of Ren et al. (2020), a dataset validated through the TemPhD method (Zhang et al., 2022), adding another 192,326 phage contigs extracted from 148,229 assemblies.

To address sequence redundancy present in both the RefSeq and TemPhD databases, we applied the CD-HIT algorithm (Fu et al., 2012; Li and Godzik, 2006) (using CD-HIT-EST with a default word size of 5). While several clustering thresholds (0.99, 0.95, 0.90) were experimented with and found to produce similar outcomes, we settled on a threshold of 0.99. This process resulted in a refined set of 40,512 distinct phage sequences, with an average length of approximately 43,356 base pairs, culminating in a total of 3.5 billion base pairs. Notably, these sequences target a wide spectrum of 660 bacterial genera. Subsequent to sequence curation, phage sequences were mapped to their respective bacterial hosts to formulate a balanced training dataset, ensuring equitable representation between phages and their hosts. This step is imperative, given the distinct distributions observed between bacterial sequences and their phage counterparts. In numerous instances, due to ambiguities in species-level identification or gaps in taxonomic data, host mapping was executed at broader taxonomic strata, predominantly at the genus level.

In our examination of bacteriophage-host associations at the genus level, several bacterial genera stood out, showcasing pronounced phage interactions. *Salmonella*, a main cause of food-related sicknesses (Popoff et al., 2004), stands out with an impressive association of 24,182 phages, spanning a cumulative length of over a billion base pairs (1,026,930,954 bp) and an average phage length of 42,467 bp. Following closely, the common gut bacterium, *Escherichia* (Tenaillon et al., 2012), is linked with 8,820 phages, accumulating a total length of 408,866,394 bp. The genus Klebsiella, notorious for its role in various infections (Paczosa and Mecsas, 2016), associates with 4,904 phages. Genera such as *Listeria* (Vázquez-Boland et al., 2011), *Staphylococcus* (Lowy, 1998), and *Pseudomonas* (Driscoll et al., 2007), each with distinct clinical significance, exhibit rich phage interactions. Notably, *Mycobacterium* (Cole et al., 1998), consisting of pathogens like the tuberculosis-causing bacterium, shows associations with 2,156 phages. Many of these bacterial genera are benign and even beneficial under normal conditions, they also include species that can cause severe diseases in humans, especially when there’s an imbalance in the body’s natural flora or when antibiotic resistance develops. Monitoring phage interactions with these bacteria offers potential pathways for therapeutic interventions and a deeper understanding of microbial ecology in human health.

Additionally, balanced databases were created, stratified by the host genus level, to mitigate the effect of underrepresented or overrepresented phages, such as Salmonella. The reverse-complement sequences were included. The final dataset encompasses a total of 660 unique bacterial genera. Undersampling was performed with a threshold of 20,027,298 bp for 25 genera, while the others were upsampled with a maximum coverage of 5x, obtaining random samples of shorter fragments from the contigs. Random segmentation and sampling were carried out as previously described. The bacterial assemblies were randomly selected from the NCBI database, prioritizing higher-quality assemblies. Many of them were not included in the pretraining dataset. Subsequently, we constructed a database with various sequence lengths: 256, 512, 1024, and 2048 bps. The train-test-validation split was executed in a 0.8, 0.1, 0.1 proportion at the phage sequence level.

For comparison with alternative methods and tools, we had to subsample our test set (*N* = 10, 000) to conduct the evaluation within a reasonable timeframe.

#### 2.4.2 Model training for phage sequence analysis

The task was formulated as binary classification, similarly to the promoters. Phage sequence classification was approached in a manner analogous to the promoter training. Given the extensive size of the dataset, preprocessing was conducted beforehand, segmenting sequences into various lengths: 256, 512, 1024, and 2048 bps. For both `mini` and `mini-c` models, the training process was partitioned into three distinct phases. An initial grid search was executed to optimize learning rates, and base models were trained for an hour. The parameter yielding the highest Matthews Correlation Coefficient (MCC) was selected. The model was then trained using segment lengths of 256 bps for half an epoch, followed by 512 bps for another half epoch, and concluding with two epochs for 1024 bps segments. The training regimen for the `mini-long` model was similar, albeit commencing with 512 bps segments, then transitioning to 1024 bps, and finally to 2048 bps segments. Model optimization employed the settings delineated previously.

### 2.5 Applied metrics

#### MCC (Matthews Correlation Coefficient)

Used for binary classifications and defined as:

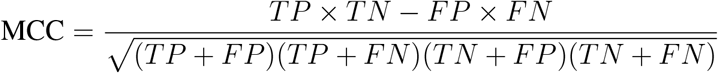

where *TP* is true positives, *TN* is true negatives, *FP* is false positives, and *FN* is false negatives. The coefficient ranges from *−*1 (total disagreement) to 1 (perfect agreement).

#### F1 Score

The harmonic mean of precision and recall, given by:

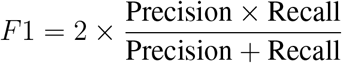

with

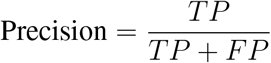

and

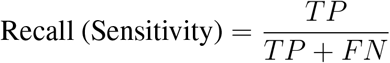

#### Accuracy

Represents the proportion of correctly predicted instances to the total, defined as:

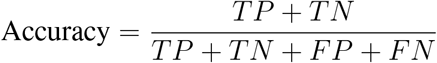

#### Sensitivity (Recall)

The proportion of actual positives correctly identified:

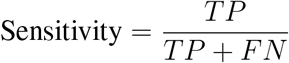

#### Specificity

The proportion of actual negatives correctly identified:

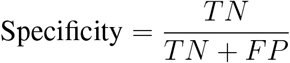

#### ROC-AUC (Receiver Operating Characteristic - Area Under Curve)

Evaluates the model’s discriminative ability between positive and negative classes. It’s the area under the ROC curve, which plots Sensitivity against 1 *−* Specificity for various thresholds.

**The silhouette score** is a measure used to calculate the goodness of a clustering algorithm. It indicates how close each sample in one cluster is to the samples in the neighboring clusters, with values ranging from -1 to 1, where a high value indicates that the sample is well matched to its own cluster and poorly matched to neighboring clusters (Rousseeuw, 1987).

Equation for the silhouette score *s*(*i*) for a single sample:

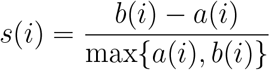

Where:

- *a*(*i*) is the average distance from the i-th sample to the other samples in the same cluster.
- *b*(*i*) is the smallest average distance from the i-th sample to samples in a different cluster, minimized over clusters.

## 3 RESULTS AND DISCUSSION

### 3.1 ProkBERT’s learned representations capture genomic structure and phylogeny

We assessed the zero-shot capabilities of our models by examining their proficiency in predicting genomic features based solely on embedding vectors, in a manner akin to Nucleotide Transformers and related methodologies. Figure 5 presents the UMAP projection of these embedded vector representations. Employing the UMAP technique, we reduced the dimensionality of genomic segments and derived embeddings. These were then evaluated using silhouette scores across the three models: `ProkBERT-mini`, `ProkBERT-mini-c`, and `ProkBERT-mini-long`.

**Figure 5.**
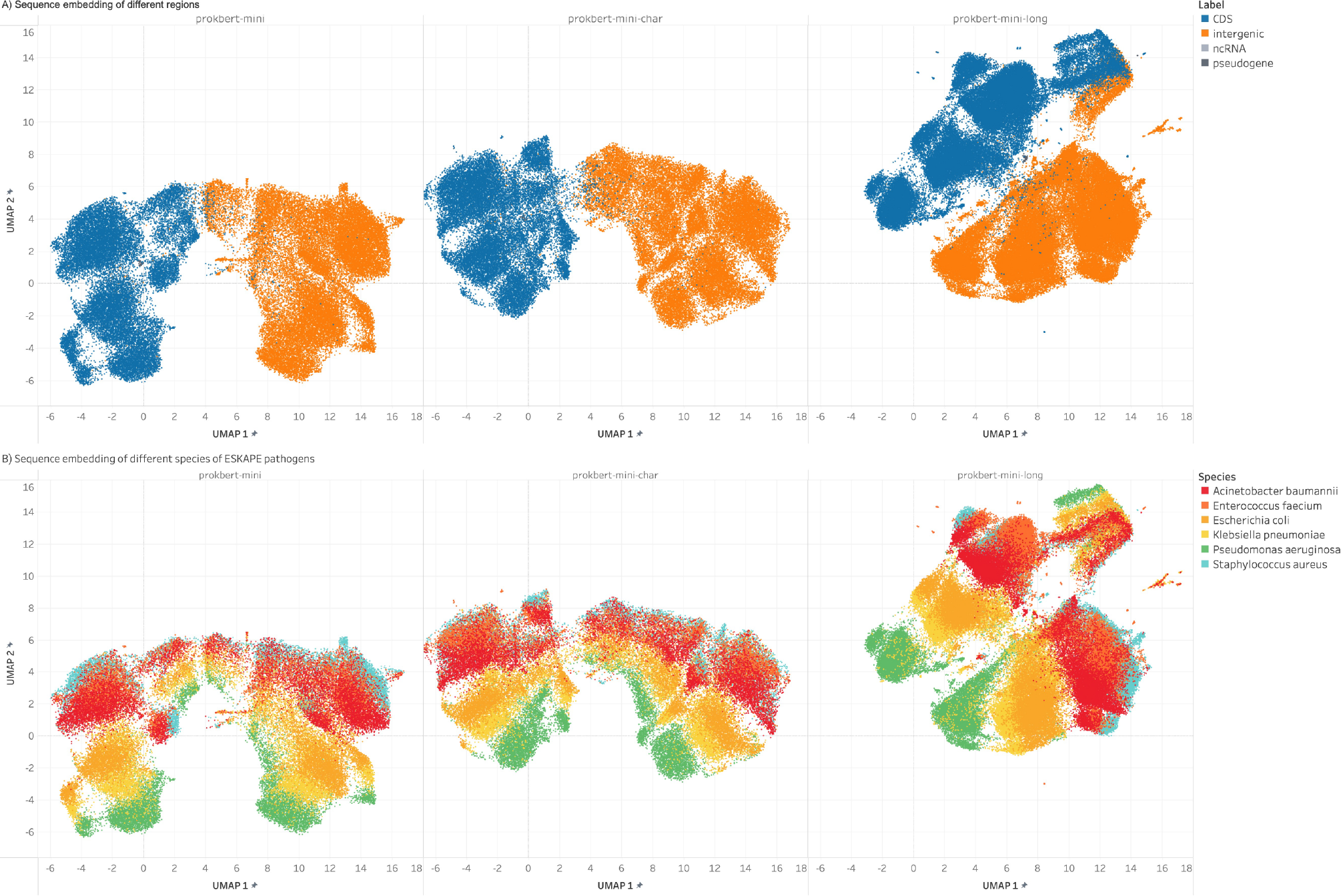
UMAP embeddings of genomic segment representations. The figure presents the two-dimensional UMAP projections of embedded vector representations for various genomic features, derived from the `ProkBERT-mini`, `ProkBERT-mini-c`, and `ProkBERT-mini-long` models. The distinct clusters highlight the models’ ability to differentiate between features such as CDS (coding sequences), intergenic regions, ncRNA, and pseudogenes, even without explicit training for feature differentiation. B) The segments are colored according to species, indicating that cluster structure reflects the phylogenic similarities.

Our primary objective was to discern if the representations of sequence segments from ESKAPE pathogens could be distinctly categorized. Indeed, Figure 5 exhibits clear delineation among known genomic features, including CDS (coding sequences), intergenic regions, ncRNA, and pseudogenes. It’s important to note that these models were not explicitly trained to differentiate these sequence features; the representations were solely derived through pretraining.

For the critical genomic comparison between ‘intergenic’ and ‘CDS’ regions, the silhouette scores obtained were 0.4925, 0.5766, and 0.3352 across the respective models, emphasizing a consistent and clear distinction between these features. Regarding non-coding RNA representations, the silhouette scores for ‘ncRNA’ vs. ‘CDS’ were 0.1537, 0.2935, and 0.2192, while for ‘ncRNA’ vs. ‘intergenic’, they were 0.1648, 0.1302, and 0.3109, further affirming the assertion that ncRNAs cluster distinctly. Pseudogenes, as anticipated, exhibited some overlap with ‘CDS’, notably in the `ProkBERT-mini` model with a score of *−* 0.0358. Yet, when compared with ‘ncRNA’, a distinct separation was observed, as evidenced by scores of 0.1630, 0.2365, and 0.1636.

This analysis aligns with biological knowledge, where pseudogenes are expected to be more similar to CDS, while ncRNAs, which have different functions and characteristics, form distinct clusters from CDS and intergenic regions. All three models appear to produce similar clustering results for the given pairs of genomic features.

The embeddings prominently display the genomic intricacies of ESKAPE pathogens. Notably, *Klebsiella pneumoniae* and *Escherichia coli*, both members of the *Enterobacteriaceae* family, exhibit close proximity in the embedding space, echoing potential genomic kinship or shared evolutionary paths. This observation is further corroborated by the low silhouette scores across the models. In contrast, species like *Pseudomonas aeruginosa* manifest as more distinct clusters, emphasizing their genetic disparities. Intriguing overlaps, such as those between differently labeled *Acinetobacter baumannii* entities, highlight potential challenges in the data or shared genomic features. Combined, the UMAP visualizations and silhouette scores provide a profound insight into species-specific genomic embeddings, revealing both shared and distinct genomic signatures.

### 3.2 ProkBERT can efficiently recover corrupted sequences

In evaluating the models’ capabilities in the masking task, we used random masking across various genomic segments, such as CDS, ncRNA, intergenic, and pseudogenes, detailed in Table 2. We measured performance with metrics like ROC-AUC and average reference rank. However, a direct model comparison presents challenges. Notably, `ProkBERT-mini-c` boasts a significantly smaller vocabulary size (9) in comparison to `ProkBERT-mini` and `ProkBERT-mini-long` (4101) This allows `ProkBERT-mini-c` to achieve higher rankings, like top3, with relative ease as it encompasses nearly the entire vocabulary (there are 4 nucleotides). Also, the local context’s representation in `ProkBERT-mini-long` is less dense, making the restoration of the masked nucleotides harder in contrast to the others.

**Table 2.** Masking performance of the ProkBERT family.

| Model | L | Avg. Ref. Rank | Avg. Top1 | Avg. Top3 | Avg. AUC |
| --- | --- | --- | --- | --- | --- |
| ProkBERT-mini | 128 | **0.9315** | **0.4497** | 0.8960 | **0.9998** |
| ProkBERT-mini-c | 128 | 0.9429 | 0.4391 | **0.8965** | 0.9504 |
| ProkBERT-mini-long | 128 | 3.9432 | 0.2164 | 0.4781 | 0.9991 |
| ProkBERT-mini | 256 | 0.8433 | 0.4848 | 0.9130 | **0.9998** |
| ProkBERT-mini-c | 256 | **0.8262** | **0.4928** | **0.9151** | 0.9565 |
| ProkBERT-mini-long | 256 | 3.5072 | 0.2470 | 0.5258 | 0.9992 |
| ProkBERT-mini | 512 | 0.8098 | 0.5056 | 0.9179 | **0.9998** |
| ProkBERT-mini-c | 512 | **0.7983** | **0.5116** | **0.9203** | 0.9580 |
| ProkBERT-mini-long | 512 | 3.3026 | 0.2669 | 0.5435 | 0.9992 |
| ProkBERT-mini | 1024 | **0.7825** | **0.5169** | **0.9227** | **0.9998** |
| ProkBERT-mini-c | 1024 | 0.7868 | 0.5128 | 0.9222 | 0.9586 |
| ProkBERT-mini-long | 1024 | 3.2082 | 0.2768 | 0.5589 | 0.9992 |

For sequences spanning 1024 nucleotides, `ProkBERT-mini` exhibited a commendable AUC of 0.9998, accompanied by top 1 and top 3 prediction accuracies of 51.69% and 92.27%, respectively. Concurrently, `ProkBERT-mini-c` achieved an AUC of 0.9586, with top 1 and top 3 accuracies at 51.28% and 92.22%. However, `ProkBERT-mini-long` reported slightly subdued figures, with an AUC of 0.9992 and top 1 and top 3 accuracies of 27.68% and 55.89%. This underscores the efficacy of the ProkBERT model family in handling genomic tasks.

A salient observation from our analysis is that a model’s prediction proficiency is intrinsically tied to the contextual size.

In our next assessment some performance nuances became evident across various genomic regions. The `prokbert-mini` model consistently stood out, especially within the Coding Sequence (CDS) and Intergenic domains. For these regions, it achieved an unmatched ROC-AUC of 0.9998. Specifically, within the CDS region, the model attained a Top1 accuracy of 50.33%, a Top3 accuracy of 91.87%, and an average reference rank of 0.811. In the Intergenic sections, these figures were 48.97%, 91.12%, and 0.843, respectively.

The `prokbert-mini-c` model also exhibited commendable performance. Within the CDS regions, this model reached a Top1 accuracy of 50.65%, a Top3 accuracy of 91.91%, and an average reference rank of 0.802. For the Intergenic regions, the metrics were 48.84%, 91.39%, and 0.839 respectively. Despite the achievements of the aforementioned models, challenges persisted across all models in the non-coding RNA (ncRNA) domains. Even the top-performing `prokbert-mini` saw its Top1 accuracy drop to 32.46%, with an average reference rank increasing to 1.202. Contrastingly, the `prokbert-mini-long`, despite its detailed design, exhibited reduced accuracies, with Top1 and Top3 accuracies of 25.18% and 52.66% across all labels, hinting at potential inefficiencies or overfitting. Collectively, these findings underscore the importance of tailored model architectures for genomic sequences and highlight the complexities of various genomic regions, laying a foundation for future targeted deep learning strategies in genomics.

### 3.3 ProkBERT performs accurately and robustly in promoter sequence recognition

Promoters play a pivotal role in gene expression by initiating the transcription process. Identifying these genomic regions is a crucial step in understanding gene regulation in bacteria.

Our first fine-tuning task centered on promoter identification. This task is primarily viewed as a binary classification, where sequences are categorized as either promoter or non-promoter. While this formulation is quite prevalent, several alternative strategies exist for addressing this issue. A notable drawback of current methods is their training on a limited set of species Chevez-Guardado and Peña-Castillo (2021), predominantly *E. coli*, with some inclusion of *Bacillus subtilis* and other key species.

As illustrated in Figure 1, our training began with a pretrained model followed by training using cross-entropy loss minimization. We evaluated the training outcomes on two datasets: a test set curated by Cassiano and Silva-Rocha (2020), and another one comprising mixed species. The models’ performance on the first dataset can be seen in Table 3.

**Table 3.** Evaluation of promoter prediction tools on *E-coli* sigma70 dataset (Transposed)

| Tool | Accuracy | MCC | Sensitivity | Specificity |
| --- | --- | --- | --- | --- |
| ProkBERT-mini | **0.87** | **0.74** | 0.90 | 0.85 |
| ProkBERT-mini-c | **0.87** | 0.73 | 0.88 | 0.85 |
| ProkBERT-mini-long | **0.87** | **0.74** | 0.89 | 0.85 |
| CNNProm | 0.72 | 0.50 | 0.95 | 0.51 |
| iPro70-FMWin | 0.76 | 0.53 | 0.84 | 0.69 |
| 70ProPred | 0.74 | 0.51 | 0.90 | 0.60 |
| iPromoter-2L | 0.64 | 0.37 | 0.94 | 0.37 |
| Multiply | 0.50 | 0.05 | 0.81 | 0.23 |
| bTSSfinder | 0.46 | -0.07 | 0.48 | 0.45 |
| BPROM | 0.56 | 0.10 | 0.20 | 0.87 |
| IBBP | 0.50 | -0.03 | 0.26 | 0.71 |
| Promotech | 0.71 | 0.43 | 0.49 | **0.90** |
| Sigma70Pred | 0.66 | 0.42 | 0.95 | 0.41 |
| iPromoter-BnCNN | 0.55 | 0.27 | **0.99** | 0.18 |
| MULTiPly | 0.54 | 0.19 | 0.92 | 0.22 |

Cassiano and Silva-Rocha (2020) had previously gauged the efficacy of several well-established tools, including BPROM (Salamov and Solovyevand, 2011), bTSSfinder (Shahmuradov et al., 2017), BacPP (de Avila e Silva et al., 2011), CNNProm (Umarov and Solovyev, 2017), IBBP (Wang et al., 2018), Virtual Footprint, iPro70-FMWin (Rahman et al., 2019), 70ProPred (He et al., 2018), iPromoter-2L (Liu et al., 2018), and MULTiPly (Zhang et al., 2019). Additionally, we incorporated newer tools like Promotech (Chevez-Guardado and Peña-Castillo, 2021) and iPromoter-BnCNN (Amin et al., 2020). These tools encompass a broad spectrum of techniques. For instance, BPROM and bTSSfinder exploit conserved and promoter element motifs. BacPP and CNNProm use neural networks for promoter predictions in *E. coli* and other bacteria based on transformed nucleotide sequences. IBBP adopts a unique image-based approach combined with logistic regression and various sequence-based features. Tools like 70ProPred, iPro70-FMWin, MULTiPly, and iPromoter-2L leverage SVM, logistic regression, and random forest methodologies, drawing upon extracted sequence features such as physicochemical properties and k-mer compositions.

The results are presented in table 3. The ProkBERT family models exhibit remarkably consistent performance across the metrics assessed. With respect to accuracy, all three tools achieve an impressive score of 0.87, marking them among the top performers in promoter prediction. This suggests that, regardless of the specific version, the underlying methodology used in the `mini` series is robust and effective.

When evaluating the balance between true and false predictions using MCC both `ProkBERT-mini` and `ProkBERT-mini-long` slightly edge out `ProkBERT-mini-c` with an MCC of 0.74 compared to 0.73 for `mini-c`. Although the difference is marginal, it might indicate subtle refinements in the `mini-long` approach.

In terms of sensitivity, which focuses on the ability to correctly identify promoters, `ProkBERT-mini` leads with a score of 0.90, closely followed by `ProkBERT-mini-long` at 0.89 and `ProkBERT-mini-c` at 0.88. This hierarchy, albeit with small differences, highlights the minute improvements achieved in the `mini` and `mini-long` versions.

Lastly, for specificity, all three versions achieve an identical score of 0.85. This uniformity underscores the consistency in their ability to correctly identify non-promoters.

In summary, while the performance across the `mini` versions is largely comparable, `ProkBERT-mini` and `ProkBERT-mini-long` display marginal advantages in certain metrics, hinting at potential refinements in these versions.

The Promotech tool demonstrates a mixed performance across the metrics. With an accuracy of 0.71, it correctly predicts the presence or absence of promoters 71% of the time. While this accuracy is lower than the top-performing tools like `ProkBERT-mini` and its variants, it is significantly better than the lower-performing tools such as Multiply and bTSSfinder.

Sensitivity for Promotech is 0.49, suggesting that it correctly identifies nearly half of the actual promoters. However, its most remarkable performance metric is its specificity, with a score of 0.90. This means Promotech is adept at identifying non-promoters, correctly classifying them 90% of the time.

Among the methods assessed, CNNProm, Sigma70Pred, iPromoter-BnCNN, and iPromoter-2L exhibit notably high sensitivity scores, signifying their pronounced ability to correctly identify promoters. Specifically, iPromoter-BnCNN leads with an exceptional sensitivity of 0.99, closely trailed by Sigma70Pred at 0.95, CNNProm at 0.95, and iPromoter-2L at 0.94. Such high sensitivity scores indicate these models’ potential in minimizing false negatives, which is crucial in applications where missing an actual promoter can have significant implications. However, it’s vital to interpret these results with caution. The high sensitivity scores, especially of iPromoter-BnCNN and Sigma70Pred, come at the expense of specificity. For instance, iPromoter-BnCNN has a notably low specificity of 0.18, implying a substantial rate of false positives. Similarly, Sigma70Pred has a specificity of 0.41. This suggests that while these models are adept at identifying promoters, they often misclassify non-promoters as promoters. An essential factor to consider in this evaluation is the training data. Given that these models were trained on *E. coli* data, their performance might be biased when evaluated on the same or closely related datasets. This lack of independence between training and testing data can lead to overly optimistic performance metrics, as the models might merely be recalling patterns they’ve already seen, rather than generalizing to novel, unseen data.

Next, we evaluated our models’ performance on a test set encompassing a broad mix of promoters, extending beyond just *E. coli*. The results are shown in Figure 6.

**Figure 6.**
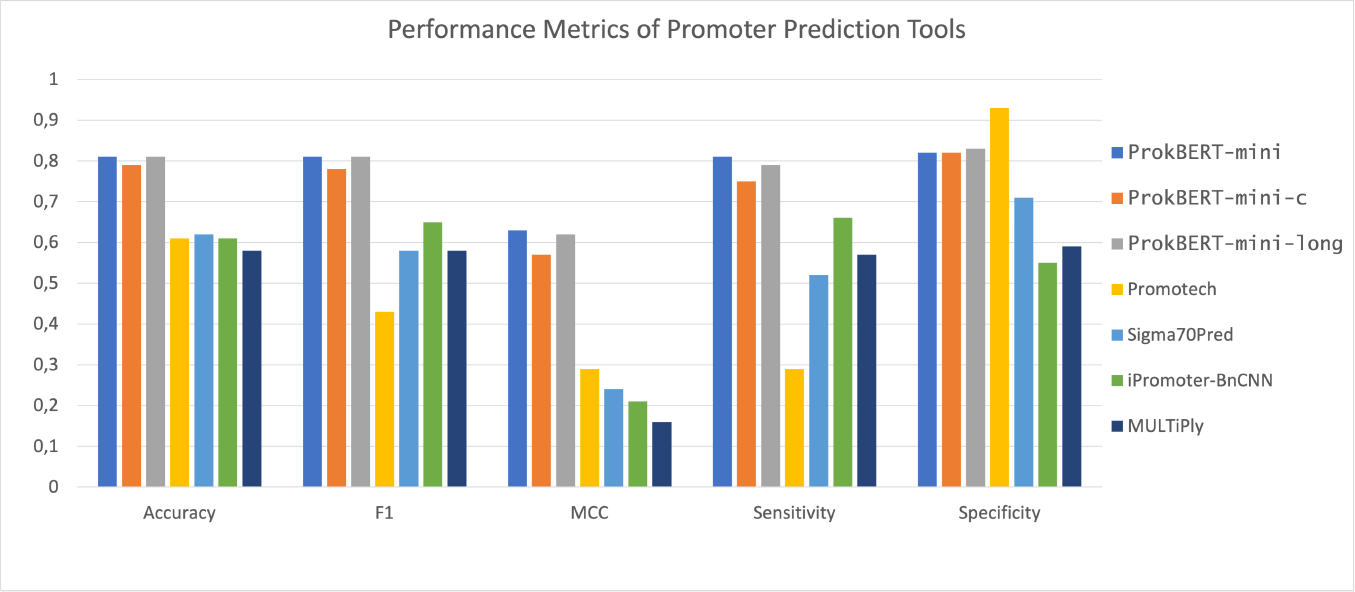
Promoter prediction performance metrics on a diverse test set. A comparative analysis of various promoter prediction tools, showcasing their performance across key metrics including accuracy, F1 score, MCC, sensitivity, and specificity. The tools evaluated include `ProkBERT-mini`, `ProkBERT-mini-c`, `ProkBERT-mini-long`, Promotech, Sigma70Pred, iPromoter-BnCNN, and MULTiPly.

The trio of tools in the ProkBERT family – `mini`, `mini-c`, and `mini-long` – consistently exhibited strong performance across the metrics analyzed. In terms of accuracy, all three achieved scores between 0.79 and 0.81, solidifying their position among leading promoter prediction tools. This uniformity in results points to a reliable methodology underlying the ProkBERT family.

Using the Matthews Correlation Coefficient (MCC) as a measure of prediction balance, `ProkBERT-mini` and `ProkBERT-mini-long` both slightly outperformed `ProkBERT-mini-c` with MCC values of 0.63 and 0.62 respectively, against the 0.57 of `mini-c`. Considering sensitivity, `ProkBERT-mini` achieved the highest score of 0.81, with `ProkBERT-mini-long` and `ProkBERT-mini-c` trailing at 0.79 and 0.75, respectively. This order reiterates the nuanced enhancements in the models. With regard to specificity, `ProkBERT-mini-long` stood out with a score of 0.83, whereas `ProkBERT-mini` and `ProkBERT-mini-c` both scored 0.82, reflecting their adeptness at accurate non-promoter classification.

Of the tools assessed, both Sigma70Pred and iPromoter-BnCNN show moderate performance in sensitivity, with iPromoter-BnCNN taking the lead at 0.66 and Sigma70Pred following at 0.52. Promotech displayed a varied metric performance. With an accuracy rate of 61%, it identifies promoters correctly in a majority of instances. Its sensitivity value of 0.29 signifies its capability to detect roughly one-third of true promoters. Yet, its high specificity of 0.93 reveals its proficiency at negating non-promoters.

Promoter prediction is an intricate task that requires a balance between sensitivity and specificity. The consistently strong performance of the ProkBERT family highlights their reliability in this domain. Yet, the selection of a tool should be made after weighing the potential implications of both false positives and negatives.

### 3.4 ProkBERT swiftly and accurately identifies phage sequences, even in challenging settings

Various tools have addressed phage sequence identification, each employing distinct strategies. These methods can be categorized into: i) homology or marker-based tools like VirSorter2 (Guo et al., 2021) and VIBRANT (Kieft et al., 2020), ii) alignment-free methods, for instance, DeepVirFinder (Ren et al., 2020) and INHERIT (Bai et al., 2022b). The first category leans on existing annotations, databases, and sequences. In contrast, alignment-free methods are less influenced by existing knowledge, offering broader applicability and greater reliability with imperfect sequence data (Wu et al., 2023). We assessed our classification accuracy against INHERIT, VirSorter2, and DeepVirFinder (Ren et al., 2020). Notably, INHERIT employs a DNABert architecture for classification, akin to ours, drawing inspiration from DNABert (Ji et al., 2021).

In genomic studies, discerning phage-related segments becomes increasingly challenging as the segment length diminishes (Guo et al., 2021). This study rigorously evaluates six distinct phage classification methodologies over a range of sequence lengths, leveraging the accuracy and MCC as primary performance metrics.

For the shortest fragments (256bp), VirSorter was unable to process the test set. Among the evaluated methods, the ProkBERT models – `mini, mini-c, mini-long` – consistently emerged as top performers across varying lengths, as depicted in Figure 7. Specifically, `ProkBERT-mini` excels with shorter sequences, achieving the highest accuracy for 256bp fragments. This high accuracy does not come at the cost of increased false positives or negatives, as evidenced by its comparable MCC values. In contrast, DeepVirFinder, ranking fifth, indicates potential optimization areas for such short sequences.

**Figure 7.**
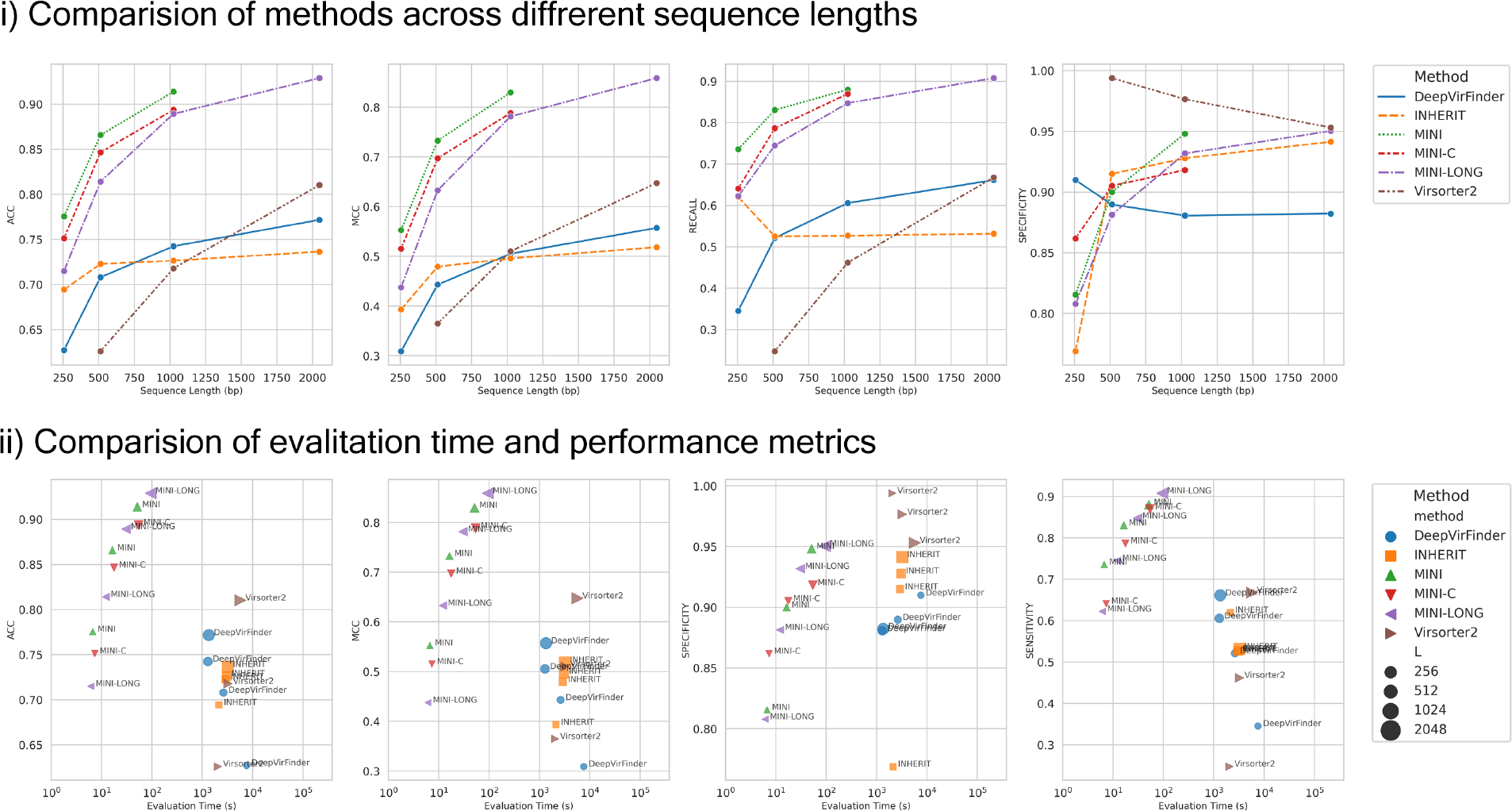
ProkBERT identifies phage sequences accurately and rapidly. i) Method comparison over varying sequence lengths based on four essential performance metrics: accuracy, MCC, sensitivity, and specificity. ii) Scatter plots illustrating the relationship between evaluation time (on a logarithmic scale) and the mentioned performance metrics. The size of each point signifies the sequence length. Evaluation time encompasses model loading, sequence preprocessing, and inference phases.

While `ProkBERT-mini` consistently ranks highest for lengths up to 1024bps, `ProkBERT-mini-c` closely follows, signifying its stability and reliability. Notably, the maximum sequence length that `ProkBERT-mini` and `ProkBERT-mini-c` can process is limited to 1024bps, introducing the specialized `ProkBERT-mini-long` for extended sequences. This model showcases its prowess with 2kb sequences, achieving an accuracy of 92.90% and an MCC of 0.859.

Virsorter2, despite initial struggles with shorter sequences, exhibits significant improvements for longer fragments. However, both DeepVirFinder and INHERIT show limited enhancements with increased sequence lengths, suggesting these methods might not capitalize on the additional information longer sequences provide as effectively as their counterparts.

In conclusion, `ProkBERT-mini` and `ProkBERT-mini-long` clearly stand out as top-performing models across various sequence lengths. While other methods may have their merits, they simply don’t match the consistency and robustness offered by the ProkBERT models.

In phage classification, sensitivity signifies the proportion of actual phage sequences that are correctly identified. Conversely, specificity represents the proportion of non-phage sequences accurately discerned. A method exhibiting high sensitivity effectively identifies most phage sequences, while high specificity indicates minimal misclassification of non-phage sequences as phage-related.

Interestingly, longer sequences tend to decrease the specificity for VirSorter2. This trend suggests that VirSorter2 might misclassify non-phage sequences more frequently as the sequence length increases. A concurrent analysis of sensitivity and specificity reveals nuances in method performance. For example, `ProkBERT-mini` consistently achieves top ranks in sensitivity but displays variable results in specificity. On the other hand, Virsorter2, despite its strong specificity, especially with extended sequences, requires enhancements in its sensitivity. Notably, several methods, including DeepVirFinder, `ProkBERT-mini`, `ProkBERT-mini-long`, and `ProkBERT-mini-c`, consistently maintain high specificity. Their narrow interquartile ranges around upper values underscore their consistent, reliable performance.

Next, we scrutinized the relationship between evaluation time and prediction performance. It’s important to note that the evaluation time encompasses not just the prediction interval but also includes sequence preprocessing and model loading durations.

The ProkBERT family shines in terms of both swiftness and efficacy. These methods, regardless of sequence length, consistently register evaluation durations under 10 seconds, making them invaluable for applications necessitating real-time predictions. Specifically, for 2kb sequences, `ProkBERT-mini-long` records a commendable accuracy of nearly 92.9%. Its Matthews Correlation Coefficient (MCC), a reliable metric of prediction prowess, stands at approximately 0.859 for the same sequence length.

In contrast, both VirSorter2 and DeepVirFinder manifest protracted evaluation phases, with the latency amplifying as sequences lengthen. Remarkably, VirSorter2 demands an evaluation span surpassing 1000 seconds for 2kb sequences. While assessing accuracy, DeepVirFinder exhibits suboptimal performance, especially with succinct sequences like 256bp, where it achieves a mere 75%. However, it’s essential to acknowledge that VirSorter2 extends beyond mere classification; it offers comprehensive annotations, a process inherently time-intensive.

In essence, the ProkBERT family epitomizes a synergy of rapidity and reliability. Concurrently, other contenders like VirSorter2, DeepVirFinder, and INHERIT unveil distinct advantages, coupled with potential avenues for refinement.

## 4 CONCLUSION

In bioinformatics, there has always been a keen interest in developing tools that can offer precise and context-sensitive interpretations of sequences. Meeting this demand, we introduced the ProkBERT model family. These innovative models benefit from transfer learning (Pan and Yang, 2009), a method showing promise in a variety of applications. A standout feature of ProkBERT is its ability to harness vast amounts of unlabeled sequence data through self-supervised learning (He et al., 2020). This approach equips ProkBERT to handle challenges like limited labeled data, a problem that has often hindered traditional models such as CNNs, RNNs, and LSTMs (LeCun et al., 2015; Cho et al., 2014). Another strength of ProkBERT is its adaptability; it performs well in different scenarios, from those with sparse data to classic supervised learning tasks (Snell et al., 2017). When we compare ProkBERT to older models that largely depend on expansive datasets, it’s clear that ProkBERT ushers in a more adaptable approach for sequence analysis in prokaryotic microbiome studies.

Our results affirm the robust generalization capabilities of the ProkBERT family. The learned representations are not only consistent but also harmonize well with established biological understanding. Specifically, the embeddings effectively delineate genomic features such as coding sequences (CDS), intergenic regions, and non-coding RNAs (ncRNA). Beyond capturing genomic attributes, the embeddings also encapsulate phylogenetic relationships. A case in point is the close proximity in the embedding space between *Klebsiella pneumoniae* and *Escherichia coli*, both belonging to the *Enterobacteriaceae* family.

We validated the versatility of the ProkBERT model family by applying it to two challenging problems: promoter sequence prediction and phage identification. Promoters play an instrumental role in transcriptomic regulation. Leveraging the transfer-learning paradigm, ProkBERT adeptly addressed the promoter prediction challenge, even when fine-tuned on multi-species datasets. This adaptability addresses a significant gap, as many conventional bioinformatics tools tend to be species-specific, often overlooking microbial diversity. In comprehensive benchmarks against prominent tools, including Multiply, Promotech, and i-Promoter2L, ProkBERT consistently outclassed both traditional machine learning and deep learning counterparts. For instance, in *E. coli* promoter recognition, it achieved an accuracy of 0.87 and an MCC of 0.74, and even in a mixed-species context, the accuracy was 0.81 with an MCC of 0.62. Additionally, our findings underscore the robustness of the training, with the`ProkBERT-mini` variant demonstrating resilience against variations in optimization parameters, such as learning rate.

Our evaluations demonstrate the prowess of ProkBERT in classifying phage sequences. Remarkably, it achieves high sensitivity and specificity even in challenging cases where available sequence information is limited. However, this exercise also highlights an inherent limitation of ProkBERT, and more broadly of transformer models: the restricted context window size. While transformer architectures are adept at capturing long-range interactions (Lin et al., 2022), they typically have a limited view of only a few kilobases.

In comparative benchmarks with varying sequence lengths, ProkBERT consistently surpassed established tools like VirSorter2 and DeepVirFinder. For instance, it attained an accuracy of 92.90% and an MCC of 0.859 in multiple benchmark studies. Intriguingly, ProkBERT even outperformed a DNA-BERT-based model, which employs a BERT architecture and vectorization strategy similar to ours.

Discussing model variants, both `ProkBERT-mini` and `ProkBERT-mini-c` have a maximum context size of 1kb, while `ProkBERT-mini-long` extends this to 2kb. Notably, `ProkBERT-mini-long` manages to use longer sequence information without compromising on prediction performance or demanding additional computational resources, thanks to the LCA tokenization strategy. Our results indicate that the local context information offered by `ProkBERT-mini-long` and `ProkBERT-mini` enhances robustness, giving them an edge over `ProkBERT-mini-c`.

ProkBERT’s superiority is not limited to prediction accuracy; it also excels in terms of inference speed. Variants such as `ProkBERT-mini`, `ProkBERT-mini-long`, and `ProkBERT-mini-c` consistently deliver outstanding performance, both in terms of evaluation speed and accuracy. Regardless of the sequence length, these models typically complete evaluations in under 10 seconds, making them exceptionally suited for real-time applications (Vaswani et al., 2017).

The vector representations generated by ProkBERT can be seamlessly integrated with traditional machine learning tools, paving the way for innovative hybrid methodologies. Being an encoder architecture, ProkBERT’s ability to produce embeddings for nucleotide sequences enables the direct incorporation of sequence information into more complex classifiers. This fusion of traditional and deep learning methods represents a promising frontier in bioinformatics. Furthermore, insights from natural language processing research suggest that the most informative representations may not always emerge from the final layer of a model (Rae et al., 2021). This underscores the need for future studies to delve deeper into the optimal layers for sequence representation extraction in bioinformatics models.

ProkBERT distinguishes itself by being both compact and powerful, embodying a blend of efficiency and accessibility. One prevailing challenge with contemporary large language models like GPT (Radford et al., 2019), BERT (Devlin et al., 2019), and T5 (Raffel et al., 2019) is their enormity. Models with hundreds of millions or even billions of parameters not only demand substantial computational resources but also complicate training and hyperparameter optimization processes. In stark contrast, ProkBERT is designed with a lean parameter count of approximately 20 million. This design choice ensures that it can comfortably fit within the memory constraints of modest GPUs. As a result, even researchers without access to high-performance computing setups or top-tier GPUs can utilize ProkBERT. Platforms like Google Colab, which offer free but limited GPU computation, become viable environments for training and evaluation tasks with ProkBERT.

In essence, our findings highlight ProkBERT’s capability to learn detailed and adaptable vector representations for sequences. These representations hold promise not only for current analytical challenges but also for emergent and unforeseen sequence classification tasks in the future. Amidst the challenges of understanding microbial communities, ProkBERT stands as a transformative tool, elucidating the complex interplay of genes and organisms in the microbiome with remarkable precision.

## AUTHOR CONTRIBUTIONS

BL: conceptualization, investigation, writing the draft and the final manuscript; BB: investigation and development; ISZN: investigation and development; NLL: investigation and writing the draft JJ: investigation and writing the final manuscript.

## ACKNOWLEDGMENTS

Thanks are due to Prof. S. Pongor (PPCU, Budapest) for help and advice. The authors extend their gratitude to all members of ML4Microbiome for their valuable discussions and feedback on this research during the ML4Microbiome meetings. The authors gratefully acknowledge the HPC RIVR consortium (www.hpc-rivr.si) and EuroHPC JU (eurohpc-ju.europa.eu) for funding this research by providing computing resources of the HPC system Vega at the Institute of Information Science (www.izum.si) as well as to HPC-KIFU Komondor.

## DATA AVAILABILITY STATEMENT

### Data Availability

The datasets generated and analyzed for this study are available on Zenodo: 10.5281/zenodo.10057832

Additionally, datasets and codes can be accessed at the following repositories:

- GitHub: https://github.com/nbrg-ppcu/prokbert
- HuggingFace: https://huggingface.co/nerualbioinfo

### Code Availability

The code associated with this study is available on GitHub: https://github.com/nbrg-ppcu/prokbert

